# The physiological effects of non-invasive brain stimulation fundamentally differ across the human cortex

**DOI:** 10.1101/639237

**Authors:** Gabriel Castrillon, Nico Sollmann, Katarzyna Kurcyus, Adeel Razi, Sandro M. Krieg, Valentin Riedl

## Abstract

Non-invasive brain stimulation reliably modulates brain activity and symptoms of neuropsychiatric disorders. However, stimulation effects substantially vary across individuals and brain regions. We combined transcranial magnetic stimulation (TMS) and functional magnetic resonance imaging (fMRI) to investigate the neuronal basis of inter-individual and inter-areal differences after TMS. We found that stimulating sensory and cognitive areas yielded fundamentally heterogeneous effects. Stimulation of occipital cortex enhanced brain-wide functional connectivity and biophysical modeling identified increased local inhibition and enhanced forward-signaling after TMS. Conversely, frontal stimulation decreased functional connectivity, associated with local disinhibition and disruptions of both feedforward and feedback connections. Finally, we identified brain-wide functional integration as a predictive marker for these heterogeneous stimulation effects in individual subjects. Together, our study suggests that modeling of local and global signaling parameters of a target area will improve the specificity of non-invasive brain stimulation for research and clinical applications.

## Introduction

Transcranial magnetic stimulation (TMS) is a unique method to non-invasively modulate human brain activity and behavior. Over recent years, TMS has steadily evolved from a scientific tool to clinical application. Repetitive TMS (rTMS) has been applied to monitor and ameliorate neurological disorders such as epilepsy (San-juan et al. 2019), pain (DosSantos et al. 2018) and stroke (McDonnell and Stinear 2017), as well as psychiatric symptoms in obsessive-compulsive disorder (Trevizol et al. 2016) and schizophrenia (Dougall et al. 2015). Only recently, the U.S. Food and Drug Administration approved rTMS as a therapeutic option for major depressive disorder (MDD) (Lefaucheur et al. 2014).

Despite its undeniable positive effect, the replicability of TMS effects varies substantially across individuals and brain regions (Diekhoff-Krebset al. 2017; Hinder et al. 2014; López-Alonso et al. 2014; Martin et al. 2003). Early studies on the motor system identified decreased cortical excitability after low-frequency (< 1Hz) stimulation (Fitzgerald, Fountain and Daskalakis 2006). This stimulation protocol has since been generalized to inhibit any cortical region and its effect has been monitored with neuroimaging methods, such as functional magnetic resonance imaging (fMRI). Numerous studies, however, have identified opposite rTMS effects across the cortex. While some groups found decreased activity after stimulation (Andoh, Matsushita and Zatorre 2015; Chen et al. 2013;Mastropasqua et al. 2014; Rahnev et al. 2013; Valchev et al. 2015; Van Der Werf et al.2010; Watanabe, Hanajima, Shirota, Tsutsumi et al. 2015), others reported increased brain activity after TMS (Cocchi, Sale, Gollo et al. 2016; Cocchi, Sale, Lord et al. 2015; Eldaief et al. 2011; Gratton et al. 2013; Mancini et al. 2017; Watanabe, Hanajima, Shirota, Ohminami et al. 2014). In summary, the inhibitory effect of low-frequency stimulation seems not to generalize from motor to other functional areas of the brain. Potentially, the heterogeneous cellular composition within target areas could shed light on region specific effects of rTMS.

Stimulating the cortical surface with TMS modulates a mixture of neuronal populations that use different neurotransmitters, perform different actions, and have different sensitivity to the stimulation (Hamada et al. 2013). Cellular data show that low-frequency stimulation modulates excitability of both gamma-aminobutyric acid (GABA-) and glutamatergic neurons and thereby has different regional effects depending on the local cellular composition (Cirillo et al. 2017). Repeated stimulation induced structural remodeling at excitatory (Vlachos et al. 2012) and inhibitory (Lenz et al. 2016) synapses and increased the expression of immediate early genes associated with synaptic plasticity (Aydin-Abidin et al. 2008) and GABA-producing enzymes (Trippe et al.2009). As the electromagnetic field of TMS spans several square centimeters of cortex, “identical stimulation protocols induce different early gene expression and not all brain regions respond equally to the magnetic stimulation” (Funke and Benali 2010). In order to increase specificity and replicability of TMS, a cross-scale theory about brain stimulation is needed that takes regional heterogeneity and underlying neurophysiology into account.

In order to meet these demands, we combined macroscopic brain imaging with generative modelling of forward (usually excitatory) and backward (usually inhibitory) connections. fMRI identifies communication between two cortical areas via functional connectivity, a measure of temporal correlation between fMRI signals (Bressler and Menon 2010). Beyond such pairwise connections, groups have recently applied graph theory methods to functional connectivity data in order to identify global metrics of a region’s functional integration into the overall brain graph (Bassett and Sporns 2017). Others have used generative modelling to explain the endogenous brain activity underlying the fMRI signal. Particularly, spectral dynamic causal modeling (DCM) rests on a biophysically plausible model of coupled (intrinsic) neuronal fluctuations in a distributed neuronal network or graph (Friston et al. 2014).

In an extensive approach, we systematically compared the cross-scale impact of identical stimulation across the human cortex. Based on different connectivity profiles for higher cognitive and early sensory regions (Gilbert and Li 2013; Riedl et al. 2016), we targeted several cortical areas and analyzed the effect of local stimulation by integrating computational modeling of cellular compartments with functional network integration on a global scale. Overall, our study revealed two major results: first, individual target identification is essential given the inter-individual variability of the macroscopic brain architecture; second, identical stimulation of sensory or cognitive regions has opposite spreading effects based on the target’s cellular composition and global network integration.

## Results

Each of the twenty-seven healthy participants underwent three counterbalanced rTMS-fMRI sessions on three different days (Fig. 1A). During each session, we identically stimulated a prefrontal (FRO), an occipital (OCC) and a temporo-parietal control (CTR) region with the aim of modulating a cognitive, sensory and a functionally heterogeneous area. We measured brain activity with resting state-fMRI before (preTMS) and immediately after stimulation (postTMS). We aimed for short transition times (mean = 5.87 min, SD = 1.1 min) between the end of stimulation and postTMS and found no significant timing differences between sessions (F(2, 44) = 3.24, p > 0.05, repeated measures ANOVA). We derived individual target regions from an online analysis of the preTMS data and loaded the coordinates into the TMS-system for continuous neuronavigation during stimulation (Fig. 1B). We then applied low-frequency (1Hz) rTMS for 20 minutes outside the MRI scanner. Twenty-three participants (twelve females, mean age = 25.74 years, SD = 3.22 years) were included in all analyses as we had to exclude two subjects who did not complete all rTMS-fMRI sessions and two subjects where we could not identify target regions during the network analysis. Please find all raw imaging data as well as analysis scripts in the online repository of OpenNEURO (see Methods for download link).

**Figure 1.**
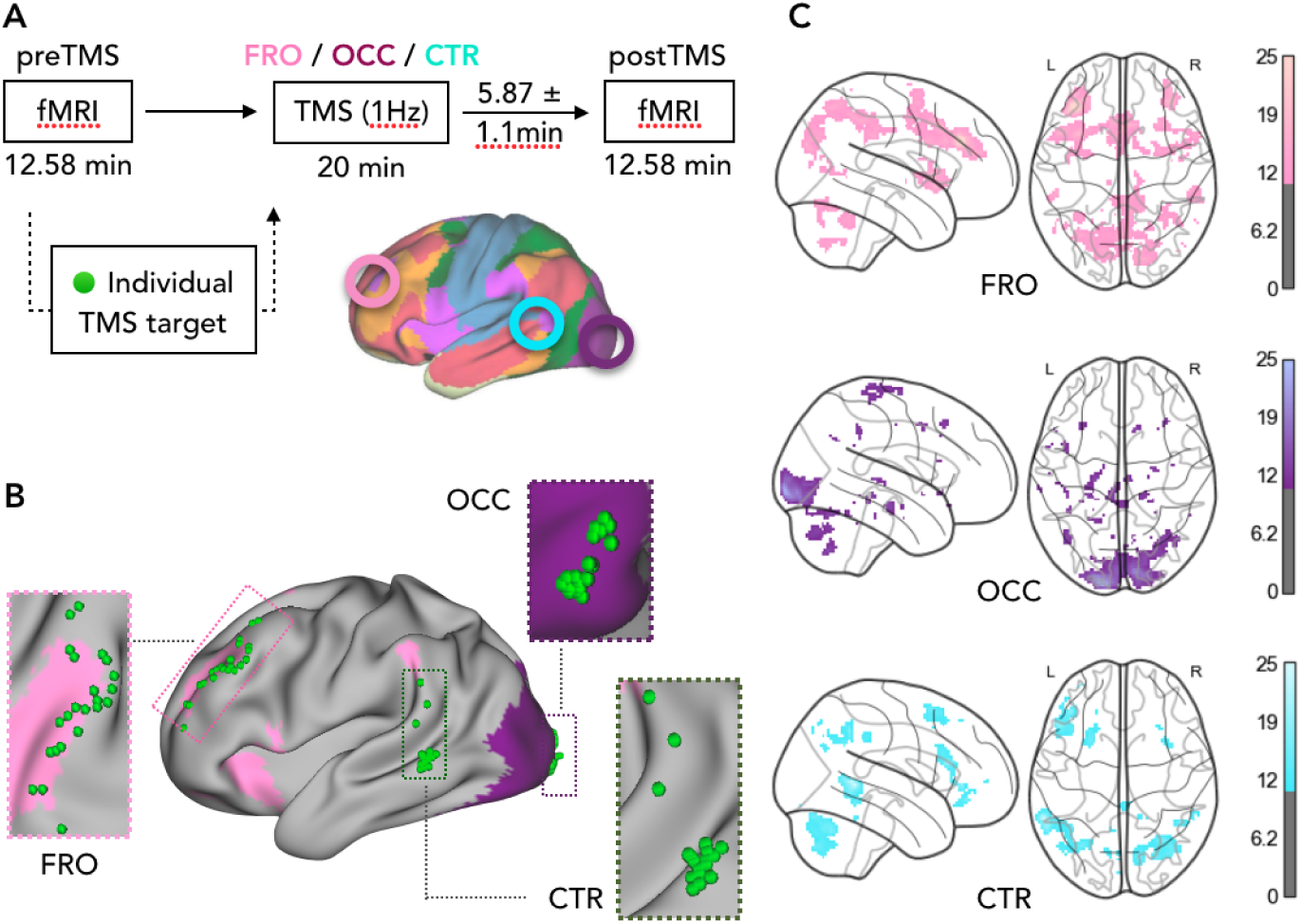
Study design. (A) Each participant underwent three counterbalanced TMS-fMRI sessions on three different days. During each session, one target region (FRO, OCC, CTR) was stimulated for 20 min with rTMS (1Hz) and we acquired resting state fMRI data during preTMS and postTMS. (B) For each subject, we derived individual target regions (green spheres) from a functional network analysis of the preTMS fMRI data. Colored overlays illustrate template networks (Yeo et al. 2011). (C) Statistical parametric maps (*p*_FWE_ < 0.05, corrected at cluster level, one-sample *t*-tests) of the group average functional connectivity of each target region during preTMS calculated from the individual TMS targets. Colorbars: *t*-values.

First, we analyzed the quality of the fMRI data in order to allow for within- and between-subject comparisons. Per session, we identified a temporal signal-to-noise ratio (SNR(*t*): mean = 6.6, SD = 1.0) and framewise displacement (FD: mean = 0.13 mm, SD = 0.03 mm) in acceptable range (Fig. S1, see Power et al. 2014) that did not differ between sessions (SNR(*t*): F(3, 66) = 1.01, *p* > 0.05; FD: F(3, 66) = 0.44, *p* > 0.05, repeated measures ANOVAs). We next validated that individually defined target areas reliably participated in the frontal (pink) and visual (violet) template networks (Yeo et al. 2011). Fig. 1C shows statistical parametric maps of voxels with significant functional connectivity with each of the target regions during preTMS (*p* < 0.05, FWE-corrected at cluster level, one-sample *t*-tests). In each subject, we therefore stimulated a sensory (visual) target, a cognitive (frontal) target, and a control target at the intersection of several networks.

### Heterogeneous spreading effects for identical stimulation protocols

We next evaluated the spreading effect of rTMS for each target region (Fig. 2). We found opposite effects after OCC-TMS and FRO-TMS with brain-wide increases (yellow voxels) and decreases (blue voxels) of functional connectivity, respectively. Fig. 2A shows separate statistical parametric maps for the *direct* impact of stimulation, e.g. increased functional connectivity of the OCC-target after OCC-TMS, but also for *indirect* effects, e.g. increased functional connectivity of the FRO-target after OCC-TMS (p_FWE_ < 0.05, corrected at cluster level, voxel-wise repeated measures ANOVAs). Interestingly, we found effects to be more spatially constrained after OCC-TMS but widespread after FRO-TMS. Consistent with stimulation targets being located in the left hemisphere, changes in functional connectivity consistently occurred in more left than right hemispheric voxels (Fig. 2A, violet bars). We found neither changes in functional connectivity for the CTR-target, nor after CTR-TMS (p_FWE_ > 0.05). Importantly, modulation effects were not selective for the target region but were spread across several template networks as illustrated by the summarized results in Fig. 2B. Overall, identical stimulation had opposite effects on functional connectivity of a sensory and cognitive region and spread to various functional networks.

**Figure 2.**
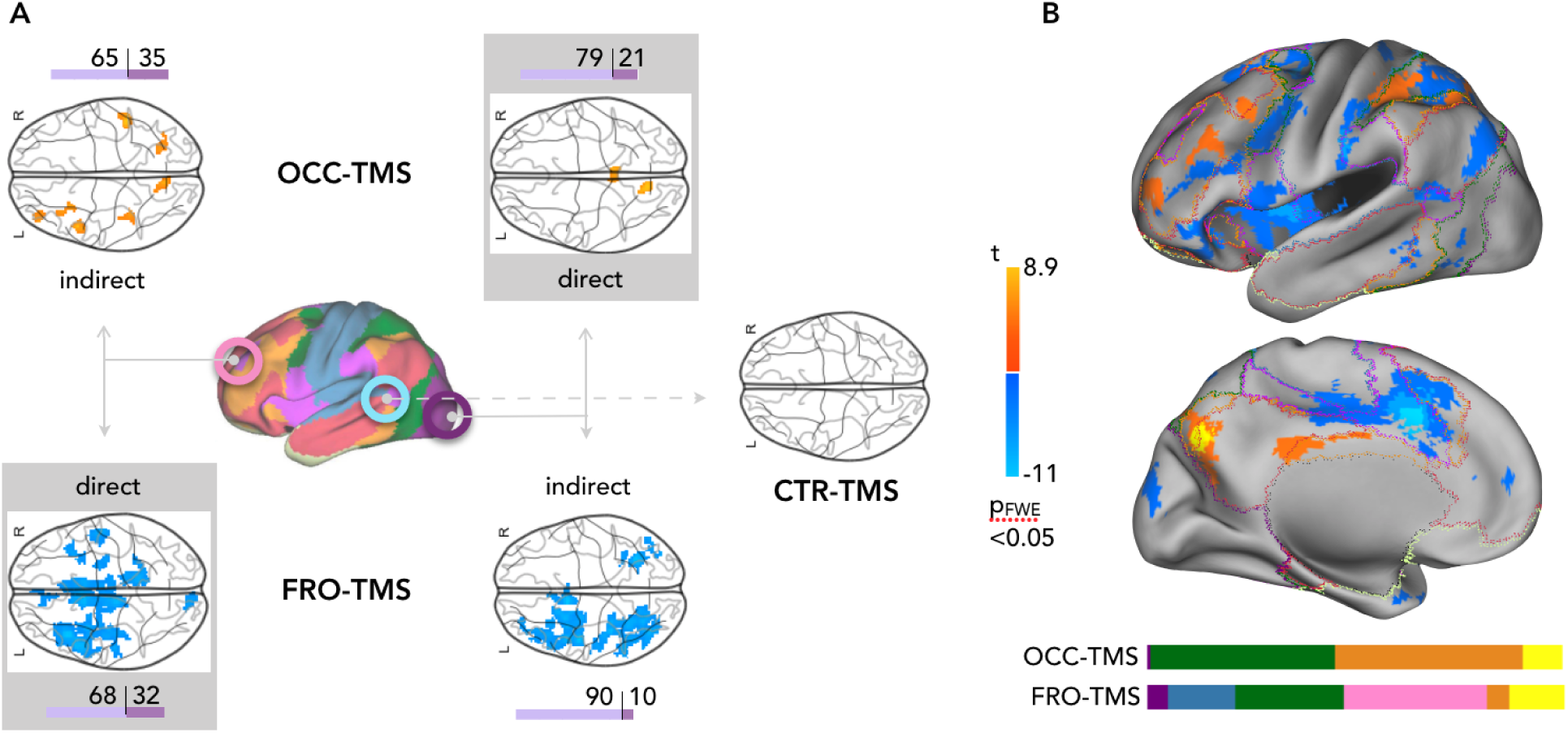
Opposite effect of TMS on brain functional connectivity. (A) Statistical parametric maps of significant changes in whole-brain functional connectivity after OCC- (upper), FRO- (lower), and CTR- (right) TMS (*p*_FWE_ < 0.05, corrected at cluster level, voxel-wise repeated measures ANOVAs). Colorbars: *t*-values. OCC-TMS increased functional connectivity of the occipital target (direct), but also of remote frontal areas (indirect). Conversely, FRO-TMS decreased overall functional connectivity to widespread cortical areas. Neither of the stimulation protocols changed functional connectivity of the CTR-target, nor did CTR-TMS. Horizontal bar plots (violet) indicate the ratio of significant voxels in left/right hemispheres. (B) Summary of whole brain-functional connectivity changes as found in (A) overlaid on the cortical surface of a standard brain. Horizontal color bars indicate the ratio of significant voxels in each of the template networks as illustrated by colored outlines on the cortical surface.

### Impact of stimulation on global functional integration

In order to investigate the stimulation effect beyond the pairwise connectivity of two regions, we next studied brain functional integration across the entire cortex. Consensus modularity analysis identified for each node, the strength of local (*z*) and global (*h*) integration within a brain graph (see Fig. 3A and Methods section). We consistently found three modules across all rTMS conditions, which were significantly more modular than comparable random networks on each of the three levels: individual FC-matrices, individual and group co-classification matrices (*p* < 0.001, permutation testing; Fig. S2). Fig. 3B shows the group co-classification matrix of preTMS as well as a topological, force-directed representation of the data. The inserts illustrate the location of the stimulation targets (grey circles) and connected nodes according to co-classification. Across subjects, the FRO-target had a significantly higher *h* than OCC-target during preTMS (*h*_*FRO*_ = 0.76 ± 013; *h*_*OCC*_ = 0.65 ± 0.19; *p* = 0.01; Wilcoxon signed rank test) which is indicative of the global integration capacity of the FRO-target between the green and blue modules.

**Figure 3.**
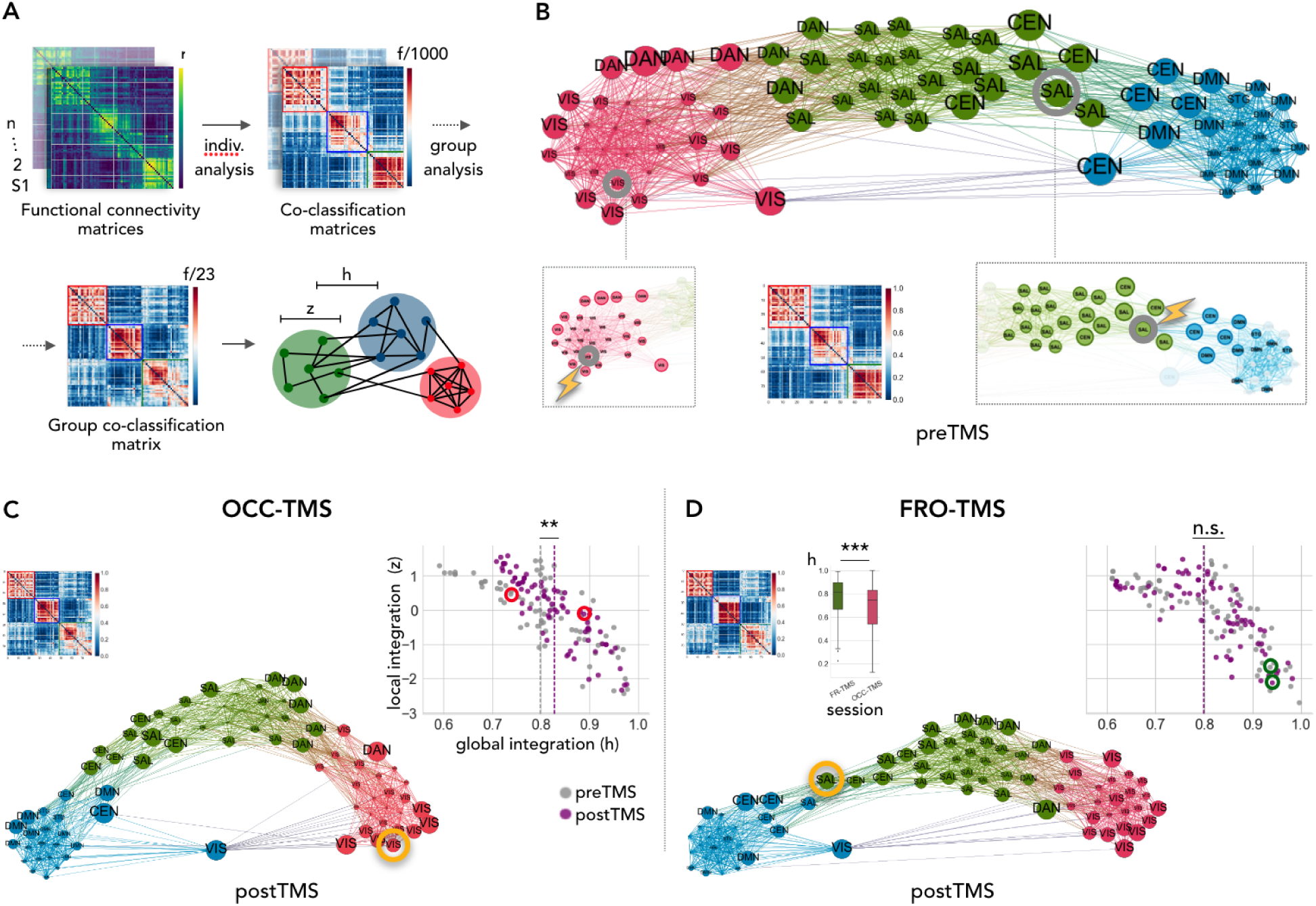
Effect of stimulation on global functional integration. (A) Overview of consensus modularity analysis on the individual and group level resulting in condition-specific parameters of local (*z*) and global (*h*) functional integration for each brain node. (B) Group average co-classification matrix and its corresponding force-directed topological representation of preTMS. Nodes with higher co-classification values are located closer to each other and node size represents *h*. Node color indicates the modular affiliation and abbreviations indicate assignment to template networks. Grey circles indicate TMS targets and inserts highlight nodes with direct functional connectivity to the target nodes. (C/D) Group average co-classification matrices and topological representations after (C) OCC-TMS and (D) FRO-TMS with target nodes (yellow circles). Scatterplots of local (*z*) vs. global (*h*) integration before (grey) and after (violet) stimulation. Note that only OCC-TMS increased global integration of the OCC-target as well as of the entire graph. ** *p* = 0.004, Wilcoxcon signed-rank test. Bar plot in (D) illustrates *h* during preTMS for all voxels that showed changes in pairwise functional connectivity. Baseline *h* was higher for voxels with spreading effects after FRO-TMS compared to OCC-TMS. *** *p* = 0.00004, Wilcoxon signed-rank test.

rTMS differentially changed the overall topology of the graph (Fig. 3C&D). Upon visual inspection, OCC-TMS arranged the modules in a more balanced, equidistant configuration (round shape of the overall graph) indicating a relative increase of exchange between red and blue modules. FRO-TMS, however, moved green and blue modules further apart indicating less consistent functional connectivity between the nodes. We also quantified the effect of stimulation on *local* and *global* integration values (scatter plots) between preTMS and postTMS. After OCC-TMS, *global* integration increased of both the OCC-target (*h*_*pre*_ = 0.65, *h*_*post*_ = 0.88, red circles Fig. 3C) as well as of the entire graph (*h*_*pre*_ = 0.80 ± 0.12, *h*_*post*_ = 0.83 ± 0.07, *p* = 0.004; Wilcoxon signed-rank test). FRO-TMS did neither impact on *global* integration of the FRO-target nor of the entire graph (*p* > 0.05, green circles Fig. 3D). The *local* integration (*z*) remained unaffected by any of the TMS interventions (p > 0.05). We also calculated *h*_*pre*_ for all voxels that showed changes in pairwise functional connectivity as illustrated in Fig. 2). Across subjects, *h*_*pre*_ was higher for all voxels that showed changes after FRO-TMS rather than for those after OCC-TMS (*p* = 0.00004, Wilcoxon signed-rank test; bar plot in Fig. 3D). This indicates that all regions affected by FRO-TMS were already more integrated at baseline which might explain why stimulation broadly affected pairwise connections but was contained on the global level. In summary, stimulation of a sensory node with low *global* integration at baseline increased brain wide interaction, possibly via direct connections to *global* integration nodes. Targeting a frontal node with initially strong *global* integration, however, had no impact on brain-wide integration, presumably as the stimulation effect was contained by the initially high level of dense connections.

### Brain functional integration as a predictive marker for spreading effects of rTMS

We next tested the discriminatory power of *global* integration for the stimulation effect at the individual subject level. We calculated the difference between pre- and postTMS *h* for a whole-brain parcellation atlas (Yeo et al. 2011) and subjected a total of N = 151 features to a random forest classifier to distinguish between OCC- and FRO-TMS sessions. Fig. 4A shows the prediction accuracy for each class (OCC-TMS = 65%; FRO-TMS = 70%; 95% CI = 53–80%), yielding an overall accuracy of 67%. Permutation testing (5000 iterations) indicated a significance level of *p* = 0.028 for the model. We replicated our classification result using a potentially more robust, linear classifier with less parameters. A support vector machine classifier yielded an overall accuracy of 65% (OCC-TMS = 65%; FRO-TMS = 65%; 95% CI = 51–78%, see Fig. S4). Finally, we explored the generalizability of our findings and calculated *global* integration parameters for the entire cortex. Fig. 4B shows the regional distribution of *h* indicating a two-part segregation of the brain. Similar to the OCC-target (violet circle), sensori-motor cortices and early integration regions have a lower *h* index (see also networks VIS, SM, DAN). Fronto-parietal cortices, covering higher cognitive networks such as SAL and CEN, have the highest *h* index, similar to our FRO-target (pink circle). In summary, *global* integration shows a two-part distribution across the cortex which might predict different response profiles of sensory and cognitive regions to low-frequency stimulation.

**Figure 4.**
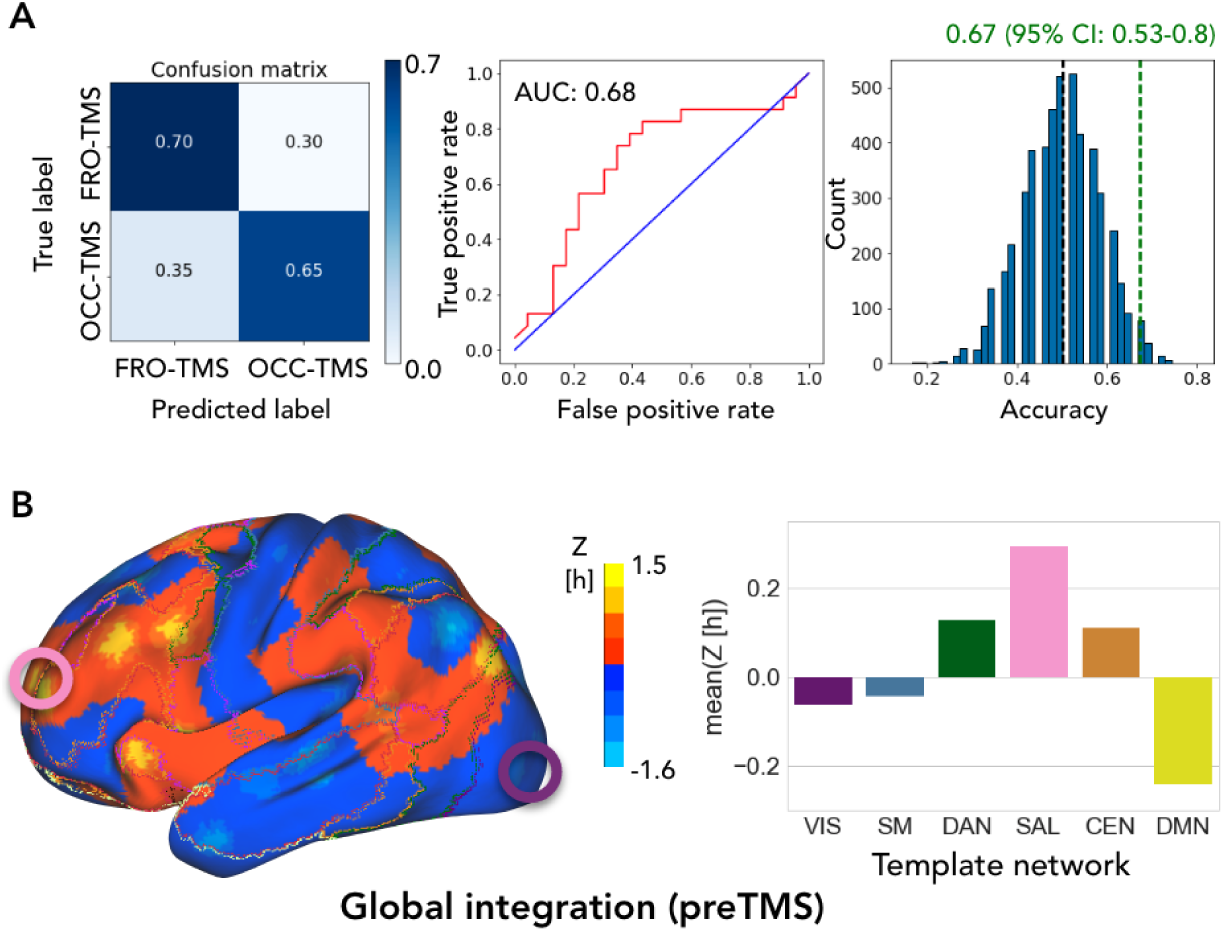
Brain functional integration is a predictive marker for spreading effects of TMS. (A) Individual classification between OCC- and FRO-TMS based on h values of a whole-brain parcellation yielded an overall prediction accuracy of 67%. (Left) Confusion matrix with the prediction accuracies for every class. (Middle) ROC curve and discrimination probability (area under the curve (AUC) of 0.68). (Right) Model (green line) significantly (*p* = 0.027, permutation testing) deviated from the null distribution (black line) after permutation testing. (B) (Left) Spatial distribution of *h* across the entire cortex, and (right) averaged across template networks.

### No effect of stimulation on the local level

Remote effects of rTMS might be simply related to local signal changes in the target region that will ultimately affect any functional connectivity measure with that region (Fox et al. 2012). We therefore analyzed three standardized imaging parameters of local fMRI signaling (Fig. 5). For each voxel in the target regions, we calculated the amplitude of low-frequency fluctuations, the regional homogeneity, and the standard deviation of the signal time-series. We found no significant effect of stimulation on any of the three parameters (*p* > 0.05, FWE corrected at the cluster level, voxel-wise repeated measures ANOVAs). Bar plots illustrate average parameters for each region and condition. This indicates that stimulation rather impacts on remote functional connectivity than on local signaling of a target region.

**Figure 5.**
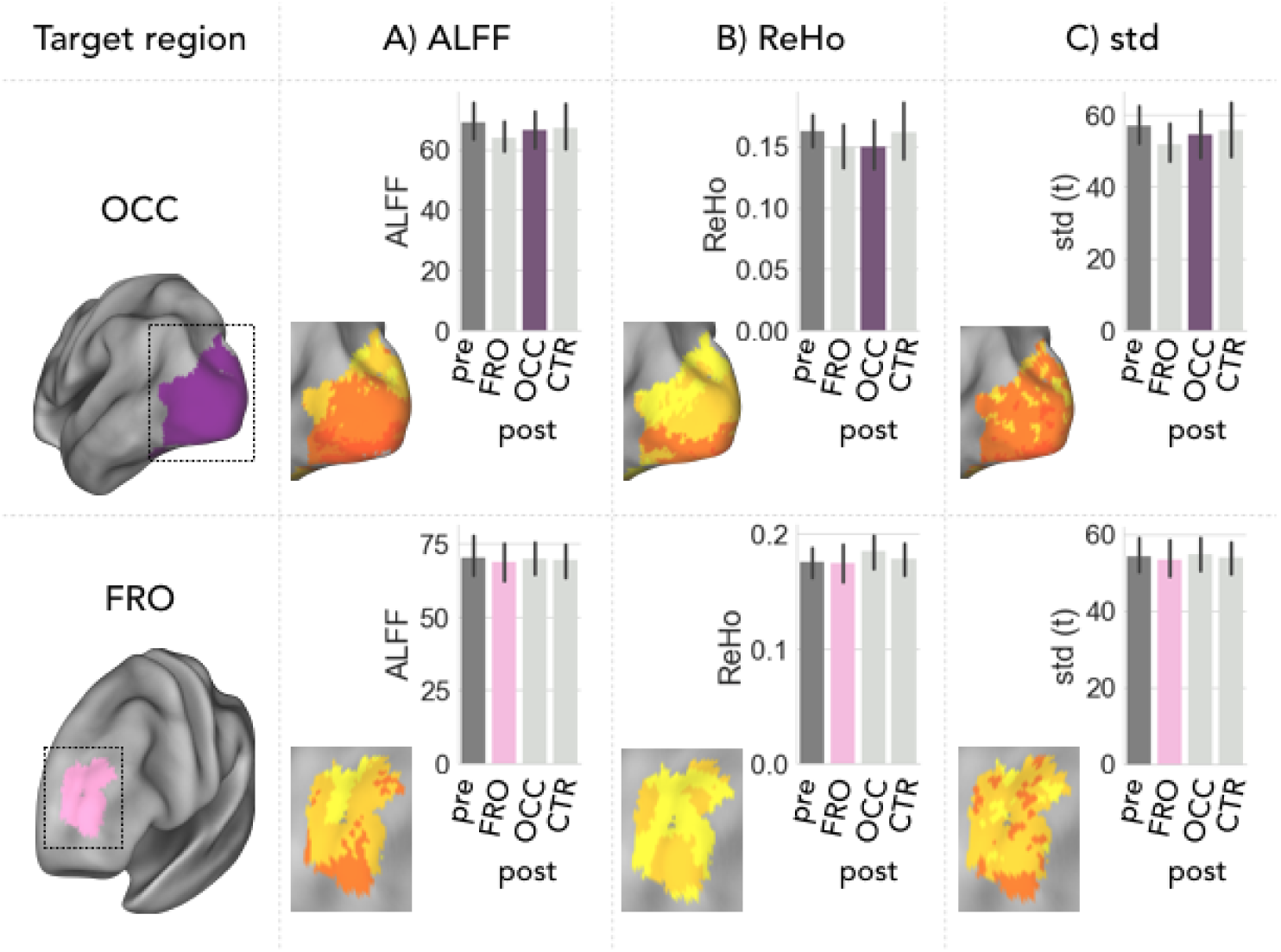
No effect of TMS on local brain activity. Bar plots indicate group average (A) amplitude of low frequency fluctuations (ALFF), B) regional homogeneity (ReHo) of local functional connectivity, and C) standard deviation (std) of the fMRI signal averaged across all voxels of each target region. PostTMS values (colored bars) after direct stimulation did not significantly differ from any other session (*p*_FWE_ > 0.05 corrected at cluster level, voxel-wise ANOVA for repeated measures). Error bars represent the 95% confidence interval of variance across subjects.

### Biophysical modeling of global stimulation effects

In a final step, we tested with generative modelling how brain stimulation differentially impacted feedforward and feedback connections among areas with functional connectivity changes. We used spectral DCM to characterize neuronal dynamics of local inhibitory and long-range excitatory connections in eight functional subnetworks of the template parcellation (R1, R2: visual- ; R5, R6: dorsal attention- ; R7, R8: salience- ; R12, R13: central executive network; Yeo et al. 2011). Fig. 6A (right) shows the group mean model of preTMS across all subjects after parametric empirical Bayes (PEB) procedures (Friston, Litvak et al. 2016). We found a balanced architecture of reciprocal connections between occipital and parietal regions. Moreover, specific feedforward (green) and feedback (yellow) connections exist along an anatomical axis of sensory, parietal integration, and frontal cognitive areas. The model also estimated inhibitory self-connections (red). Fig. 6B shows group differences in directional connectivity after OCC-TMS. Occipital stimulation increased local self-inhibition in the occipital cortex (R1: +0.13) and shifted the balance of directional connectivity to feedforward signaling towards parietal and frontal cortices (decreased feedback onto regions R1: −0.14 [from R13], R6: −0.10 [from R7 & R13], increased feedforward from R2: 0.10 [to R6]). Conversely, FRO-TMS caused global uncoupling among all regions. Fig. 6C shows decreased local self-inhibition in frontal cortex (R8: −0.10) and uncoupling of 11 out of 16 directional pathways present during preTMS (dotted lines). Stimulation equally affected feedforward and feedback connections. This analysis extends the pattern of functional connectivity changes with a generative model about asymmetric cortical hierarchies in terms of feedforward and feedback connections. Stimulation of occipital regions fosters forward signaling of regions along the visual stream while FRO-TMS leads to uncoupling of bidirectional pathways even in remote areas.

**Figure 6.**
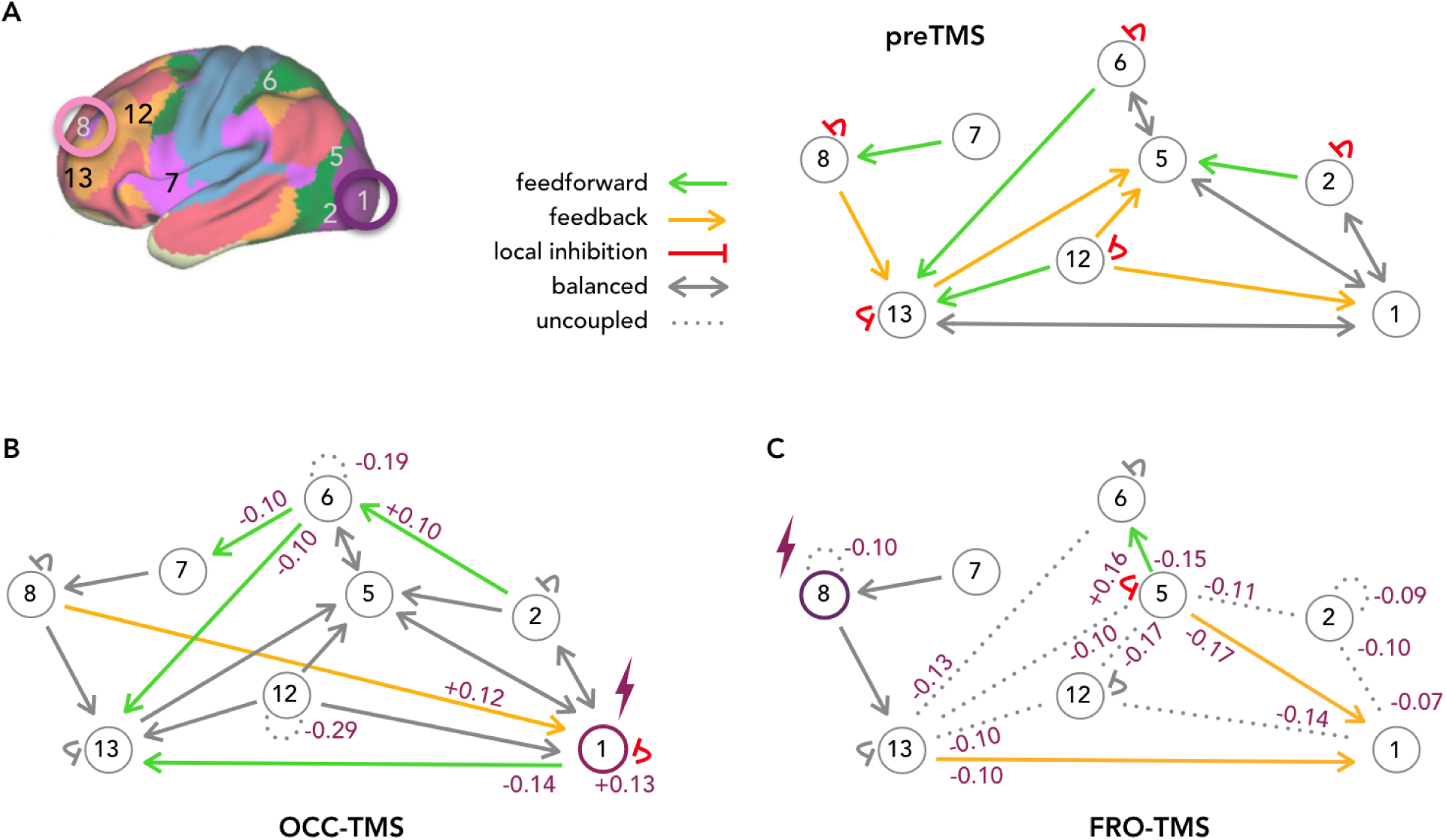
Biophysical modeling of the stimulation effect on directional signaling. (A) We modeled directional signaling between regions with functional connectivity changes using spectral DCM. The model incorporates local self inhibition (red), excitatory feedforward (green) and inhibitory feedback (yellow) signaling. Connectivity schemas illustrate directional signaling along an anterior-posterior (left-to-right) axis of the human cortex. During preTMS, the model indicated specific feedforward and feedback, as well as balanced (grey) connectivity pathways. B) OCC-TMS strengthened feedforward signaling of occipital (R1, R2) and parietal (R6) regions while other connections remained in place (grey arrows). C) FRO-TMS uncoupled 11 of the 16 directional pathways (dotted lines) equally affecting feedforward and feedback connections. Violet numbers indicate changes in parameter estimates after parametric empirical Bayes procedures with a posterior probability of > 95%.

## Discussion

Neuromodulation with TMS has attracted attention as it potentially offers an elegant way to non-invasively treat aberrant brain activity in neuropsychiatric disorders. Reports about low outcome and replicability, however, have partly compromised this method (Lage et al. 2016; Martin et al. 2003; Wilson et al. 2018). We hypothesized that the key assumption about a frequency-dependent effect of rTMS does not generalize across the entire cortex and could be responsible for inconsistent reports. We found that identical stimulation had fundamentally different effects on macroscopic network signaling for sensory and cognitive areas. Generative modeling suggests that a heterogeneous cellular composition of local inhibition and remote excitation is responsible for these different response profiles. Moreover, we identified functional integration as a reliable parameter to predict a region’s response to low-frequency stimulation. Our findings provide experimental evidence for a heterogeneous signaling architecture in the human brain. Moreover, we provide a theoretical and practical framework to correctly target and increase specificity of brain stimulation in humans.

The most striking observation from our study is that low-frequency stimulation to different areas had opposite modulatory effects on that regions’ remote communication. Until now, low-frequency stimulation is assumed to decrease neuronal excitability independent of the target location (Fitzgerald, Fountain and Daskalakis 2006). This is surprising as some studies have indicated diverse effects after stimulating different parts of the visual system (Cocchi, Sale, Gollo et al. 2016; Ruff et al. 2006) and several groups have repeatedly reported increased rather than decreased activity after low-frequency rTMS of various cortical regions (Cocchi, Sale, Lordet al. 2015; Eldaief et al. 2011; Gratton et al. 2013; Mancini et al. 2017; Watanabe, Hanajima, Shirota, Ohminami et al. 2014). Yet, a systematic comparison of local and global effects of stimulation to different functional areas has been missing so far. Our results suggest that the effect of modulation varies across the cortical surface and is rather determined by the extent of functional integration of a target area than by the frequency range of the stimulation protocol.

The most consistent result we identified across regions is that stimulation broadly spreads beyond the target area and associated functional networks. This remote impact is consistent with studies that used TMS to modulate functional connectivity. However, reports have focused on the stimulation effect within a certain functional system, such as the visual (Rahnev et al. 2013), sensory (Andoh, Matsushita and Zatorre 2015; Valchev et al. 2015), motor (Watanabe, Hanajima, Shirota, Tsutsumi et al. 2015), or default mode (Eldaief et al. 2011; Van Der Werfet al. 2010) network. Such findings tend to convey the impression that TMS is capable of modulating a specific network in isolation. Our whole-brain approach revealed effects beyond the functional system covering the target area. Occipital stimulation had no effect on the visual system, but spread to parietal and frontal regions along the entire dorsal visual stream (Mishkin, Ungerleider and Macko 1983). This is in line with anterograde tracing data, which revealed that the majority of neurons in V1 mainly broadcast to cortical and subcortical regions outside the visual system (Han et al. 2018). Conversely, frontal stimulation decreased functional connectivity to all major parts of the salience network (covering the frontal target) and to template networks across the entire cortex. This result reflects the high level of distributed connectivity which is inherent to the frontal cortex as identified on the micro- (Modha and Singh 2010) and macro- (Power, Cohen et al. 2011; Van den Heuvel and Sporns 2013) scale. In summary, we identified a dichotomy of specific effects after sensory, and broad effects after frontal stimulation, which reflects an established signaling hierarchy of divergence and convergence in the human cortex based on computational modeling (Man et al.2013) and tract tracing studies (Modha and Singh 2010). Therefore, rTMS seems to be a promising tool to identify, and also modulate, particular functional pathways. However, it is important to note that spreading effects will not be confined to the network of interest alone.

We propose that functional integration is a suitable marker to predict specific response patterns to stimulation. Functional integration is a simple measure illustrating a region’s connectivity profile within the whole-brain graph. Based on the integration parameters of all cortical nodes, we successfully classified whether occipital or frontal stimulation was applied to the individual subject. Interestingly, functional integration consistently distinguished among sensory and higher cognitive systems in general. This suggests that sensory and cognitive areas will respond similarly to stimulation as did the occipital and frontal cortex, respectively. The question then arises: How to integrate our results of pairwise functional connectivity and global integration? Occipital stimulation selectively strengthened functional connectivity with regions distributed along the dorsal visual stream. This suggests that occipital stimulation prepares the entire brain network for visual input and is in line with animal data about task-dependent activation of long-range projection neurons in sensory cortices (Chen, J. L. et al. 2013). Our observation that frontal stimulation did not impact global integration is astonishing in light of the widespread decline of pairwise functional connectivity. However, such a compensatory effect has been predicted by virtual lesion modeling of brain graphs. These studies revealed that highly integrated nodes are more resilient against massive loss of individual connections (Avena-Koenigsberger et al. 2017). In summary, rTMS effects manifest with varying, and even partly opposing characteristics; yet they can be consistently interpreted across different scales.

Generative modeling using spectral DCM revealed that distinct stimulation effects potentially relate to a heterogeneous cellular composition in target areas. rTMS differentially modulated short-range inhibitory and long-range excitatory signaling in occipital and frontal areas. The model response to occipital stimulation was similar to a reported shift in the excitation/inhibition balance observed during visual processing (Liu et al. 2011). For example, Haider et al. showed that local inhibition dominates excitation in amplitude and over time during awake visual processing (Haider, Häusser and Carandini 2013). Information processing then propagates along the functional hierarchy of the dorsal visual stream via excitatory forward signaling (Mishkin, Ungerleider and Macko 1983). Frontal stimulation, on the other hand, led to uncoupling of various feedback and feedforward pathways reaching down to sensory areas. This is in line with imaging data about diverse connections of salience regions with cognitive and sensory areas (Power, Cohen et al. 2011). Moreover, our results on the macroscopic level converge with predominantly reciprocal connections among long-range pyramidal cells (Harris and Mrsic-Flogel 2013) and recent tract tracing studies in primates revealing an underrepresented network of long-range feedback connections of frontal onto sensory cortices (Markov et al. 2013).

Overall, our study could provide guidance for future applications of rTMS in clinical settings. Specifically, our design addressed repeated critique about the regional specificity of rTMS by systematically studying different cortical areas (Polanía, Nitsche and Ruff 2018): We used i) electric-field neuronavigation to identify functional target regions in each individual and, ii) neuroimaging to study the effects of stimulation beyond the stimulated network. iii) We included stimulation of a control region (in contrast to sham stimulation) to test the functional specificity of our target areas. iv) We finally integrated macroscopic findings on the network level with a generative model to propose cellular mechanisms related to electromagnetic stimulation. It is important to note that stimulation of a cognitive network spreads to the entire brain. The strong interconnectedness of a cognitive hub might yet compensate for the local impact and diminish the effect of stimulation. Repeated applications are therefore necessary to achieve a prolonged effect in areas with high integration capacity (Lefaucheur et al. 2014). On the other hand, a number of studies have specifically identified altered functional connectivity of the default mode network in neuropsychiatric disorders (Whitfield-Gabrieli and Ford 2012). The default mode network is among the densest connected network of the brain, yet with less diverse integration (Power, Cohen et al. 2011; Van den Heuvel and Sporns 2013). Its global integration profile therefore indicates the default mode network as an interesting target for efficient and rather specific modulation with non-invasive brain stimulation.

## Acknowledgements

This study was conducted with financial support to V.R. from the Deutsche Forschungsgemeinschaft (DFG, grant agreement No: 273427765) and the European Research Council (ERC) under the European Union’s Horizon 2020 research and innovation programme (grant agreement No: 759659). G.C. received a scholarship from the Colombian Government (COLCIENCIAS scholarship, grant agreement No: 646, 2014). A.R. is funded by the Australian Research Council Discovery Early Career Research Award Fellowship (DE170100128) and the Wellcome Trust.

## Author contributions

Conceptualization: V.R.; Methodology: G.C., A.R.; Formal Analysis: G.C.; Investigation: G.C., N.S., and K.K.; Resources: S.M.K.; Data Curation: G.C. and V.R.; Writing - Original Draft: G.C. and V.R.; Writing - Review & Editing: N.S., A.R., S.M.K.; Visualization: G.C. and V.R.; Supervision: V.R.

## Methods

### Participants

Twenty-seven healthy participants (fourteen females, mean age = 25.56 years, SD = 3.01 years), right-handed and without any psychiatric condition, were informed of the objectives and potential risks of the study and signed a written consent inform. The study was approved by the local institutional review board and was conducted in accordance with the Declaration of Helsinki.

### Brain stimulation

rTMS was delivered using an electric-field-navigated Nexstim eXimia system (version 4.3; Nexstim Plc, Helsinki, Finland) and a biphasic figure-of-eight stimulation coil. Before any rTMS session, the neuronavigation system was set up by co-registering the participants’ head to their structural MRI data (T1-weighted 3D-TFE sequence), allowing to continuously track the coil position with respect to the individual target region via infrared cameras. We identified individual resting motor thresholds (rMT) according to the maximum likelihood algorithm by mapping the cortical representation of the right abductor pollicis brevis muscle using surface muscle electrodes [Neuroline 720; Ambu, Ballerup, Denmark] and an integrated electromyography device (Awiszus 2003; Rossini et al. 2015; Sollmann et al. 2016). rTMS target regions were individually identified during a network analysis (see below) of the preTMS fMRI data and then overlaid onto structural images to guide the stimulation. Low-frequency rTMS with a frequency of 1 Hz and a stimulation intensity of 100% of the individual rMT (mean rMT = 34.4 %, SD = 7.5 %), was applied for 20 minutes to each of the target regions (1200 rTMS pulses in total) in a magnetically shielded room next to the MRI scanner. During stimulation, the coil was positioned perpendicular in relation to the skull surface, and an anterior-posterior orientation of the induced electric field was maintained using an adjustable coil holder during stimulation application.

### Image acquisition

MRI data were acquired on a 3T Philips Ingenia MRI scanner using the body coil for transmission and the 32 channel head coil for signal reception (Philips Healthcare, Best, The Netherlands). We acquired multiband fMRI data during each pre- and postTMS condition (40 slices; multiband factor, MB=2, SENSE factor, s=2; repetition time, TR=1250ms; echo time, TE=30ms; flip angle, FA=70°; field of view, FOV=192×192mm; matrix size=64×64; voxel size=3×3×3mm). Each fMRI run lasted for 12.35 minutes, during which 600 functional volumes were acquired. Additionally, we acquired a T1-weighted 3D-TFE during each session (170 slices; repetition time, TR=9ms; echo time, TE=3.98ms; flip angle, FA=8°; field of view, FOV=256×256mm; matrix size=256×256; voxel size=1×1×1mm). During fMRI acquisition, the scanner room was dimmed and the participants were asked to stay awake with their eyes open. This was ensured by monitoring their eyes with an MR-compatible infrared camera (12M, MRC Systems, Heidelberg, Germany) attached to the coil.

### Image data processing and analysis

We deposited the raw imaging data and analysis scripts in the online repository of OpenNEURO (https://openneuro.org/datasets/ds001927) to allow for replication and further analyses. We performed pre-processing of the structural and functional MRI data using version 0.392 of the configurable pipeline for the analysis of connectomes (C-PAC, Craddock et al. 2013).

### Pre-processing structural images

The structural images were skull-stripped using AFNI-3dSkullStrip (Cox 1996), segmented into three tissue types using FSL-FAST (Zhang, Brady and S. Smith 2001) and constrained into the individual participant tissue segmentations from standard space provided by FSL. They were then normalized to the Montreal Neurological Institute (MNI) 152 stereotactic space (2 mm3 isotropic) with linear and non-linear registrations using FSL-FLIRT (Jenkinson et al. 2002) and FSL-FNIRT (Andersson, Jenkinson, S. Smith et al. 2007), respectively.

### Pre-processing functional images

The functional images of each run were realigned, motion corrected to the average image using AFNI-3dvolreg, and then skull-stripped using AFNI-3dAutomask. Subsequently, the global mean intensity was normalized to 10,000, the nuisance signal was regressed, and the signal was bandpass filtered (0.01 - 0.1 Hz). Furthermore, the pre-processed images were registered to the structural space with FSL-FLIRT using a linear transformation based on the white matter boundary information derived from the prior white matter tissue segmentation from FSL-FAST. The nuisance signal regression step modeled the scanner drift using quadratic and linear detrending, while the physiological noise was modeled using the 5 principal components with the highest variance from a decomposition of white matter and CSF voxel time-series (CompCor, Behzadi et al. 2007), which were derived from the prior tissue segmentations transformed from anatomical to functional space. Furthermore, the head motion was modeled using the 24 regressors derived from the parameters estimated during motion realignment based on the Friston 24-Parameters (Friston, Williams et al. 1996), the six head motion parameters and their 12 corresponding squared values. If not states otherwise below, fMRI analyses were all performed in individual space. Only results were later transformed into MNI space for group statistics by applying each individual’s MNI-transformation parameters of the structural image to the results maps.

### Identification of individual target regions

We exported the fMRI data to an external computer while the remaining MRI-protocol of preTMS was still running. We ran an independent component analysis (ICA) of the preTMS data using FSL-MELODIC (Beckmann and Smith 2004) and decomposed the data into 17 spatial components. We then calculated the spatial cross-correlation between each individual ICA map and the following target networks from a template set of 17 networks by Yeo et al. (Yeo et al. 2011): for FRO: IC_08; for OCC: IC_01; for CTR: IC_14). From each of the matching ICA maps we then extracted the left hemispheric target areas covering the dorsoanterior prefrontal cortex (as FRO target), the occipital pole (as OCC target), and of the superior temporal gyrus (as CTR target), backprojected them into individual space and transferred them as TMS targets onto the navigated TMS system for immediate stimulation.

### Functional connectivity analysis

Functional connectivity is calculated as the pairwise Pearson’s correlation between the average time-series of voxels in a seed region and the time-series of all other gray matter voxels. In order to obtain least distorted connectivity patterns for each subject, we used individual TMS target coordinates as seed regions (5mm radius spheres) and calculated functional connectivity patterns in individual subject space. The preTMS functional connectivity patterns of each subject were averaged to achieve a robust pattern of baseline functional connectivity for each individual. Before we applied spatial statistics on the group level, individual functional connectivity maps were registered to the standard MNI space, Z-score transformed and spatial smoothed with a Gaussian kernel with an FWHM of 6 mm^3^.

### Consensus modularity analysis

Consensus modularity analysis, a graph theoretic method, identifies a unified partitioning of several graphs into non-overlapping clusters of nodes, i.e. modules (Dwyer et al. 2014; Fornito et al. 2012; Lancichinetti and Fortunato 2012). This analysis was implemented with MATLAB 2015b and the Brain Connectivity Toolbox (Rubinov and Sporns 2010, https://sites.google.com/site/bctnet). We visualized the modularity decomposition using the force-directed layout representation ForceAtlas2 (Gephi, Jacomy et al. 2014). First, we calculated an unthresholded FC-matrix on the individual level (per condition × session) using an independent parcellation atlas previously used for brain graph analysis (Power, Cohen et al. 2011). From this atlas, we selected those nodes (5mm radius spheres) that shared the majority of their voxels with both our group mask of grey matter as well as with any of the cognitive template networks. For each of the remaining 79 nodes, we extracted the average time-series and created a cross-correlation matrix based on the pairwise Pearson’s correlation coefficient. We zeroed negative correlation values, as well as values between nodes located within a 20 mm radius (Power, Cohen et al. 2011), and applied a Fisher *z*-transformation. Next, we ran a consensus modularity analysis on the individual level. We iteratively (1000x) applied the Louvain-algorithm for community-detection (Blondel et al. 2008) and created an individual co-classification matrix representing the frequency with which nodes were co-classified into the same module. Following recommendations in prior reports (Lancichinetti and Fortunato 2012), we chose a threshold of t=04 for the consensus partition and iterated the process 100 times. We also present results for a range of τ-values in Fig. S3 and validated the consistency of our finding with more repetitions (1000 times). Finally, we subjected individual co-classification matrices to a second-level consensus modularity analysis using identical parameters as above. The output of this analysis were group co-classification matrices per condition and two modularity parameters, classification consistency (*z*) and classification diversity (*h*), that illustrate a node’s local and global integration within the overall graph (Dwyer et al. 2014; Fornito et al. 2012).

### Local signal analysis

We calculated the amplitude of low-frequency fluctuations (ALFF, Zou et al. 2008), regional homogeneity (ReHo, Zang et al. 2004) and the standard deviation (std) of the signal time-series for each voxel within the stimulation region. For the analysis we used the preprocessed fMRI data with the following exceptions: The ALFF maps were calculated by computing the total power within the frequency range between 0.01 and 0.1 Hz of the non-filtered data, whereas the ReHo maps were calculated based on Kendall’s correlation between each voxel’s time-series and the time-series of the 27 voxels in contact with that voxel. Both measures were calculated in original space and subsequently transformed into the standard MNI space and spatially smoothed by a Gaussian kernel with a Full-Width Half Maximum (FWHM) of 6 mm^3^.

### Spectral DCM

The DCM analyses were conducted using DCM12 implemented in the SPM12 (revision 7279, www.fil.ion.ucl.ac.uk/spm). First, we projected voxel patterns of significant functional connectivity changes onto the template networks and extracted average fMRI signal time-series for eight subnetworks covering affected voxels (R1, R2: visual -; R5, R6: dorsal attention -; R7, R8: salience -; R12, R13: central executive network; Yeo et al. 2011). Then, we specified a DCM for each participant and a fully connected DCM model was created to compare all possible nested models of network interactions (Friston, Litvak et al. 2016). The model was estimated using spectral DCM, which fits the complex cross-spectral density using a power-law model of endogenous neuronal fluctuations (Friston, Kahan et al. 2014; Razi et al. 2015).

### Statistical analysis

#### Mass-univariate voxel analysis

We performed voxel-wise group statistics applying one-sample *t*-tests or one-way repeated measures ANOVAs to the parameter maps of functional connectivity, ALFF, ReHo, and std using SPM12 (http://www.fil.ion.ucl.ac.uk/spm). We configured a ‘flexible factorial design’ in SPM12 with ‘subjects’ as between-subject factor and ‘condition’ (levels: preTMS, FRO-TMS, VIS-TMS, CTR-TMS) as within-subject factor. Statistical testing was limited to voxels within an average gray matter mask derived from all participants. We corrected for multiple testing by applying a cluster-defining height-threshold of *p* = 0.001 and a cluster-extent threshold of *p* < 0.05, FWE-corrected.

#### Modularity analysis

We validated the modular decomposition using permutation testing on three levels: (i) the individual unthresholded FC-matrix and (ii) the co-classification matrix of each participant, as well as (iii) the co-classification matrix at the group level (Dwyer et al. 2014). On the individual level, we created random matrices matching the empirical matrices in degree, strength, and sign distribution per participant, and applied the identical modularity decomposition as described above. This process was repeated 5000 times, generating a null distribution of median *Q* values against which we compared the magnitude of the observed sample median *Q* per condition (Dwyer et al. 2014; Fornito, Zalesky and Bullmore 2016; Rubinov and Sporns 2011). A Wilcoxon signed-rank test (*p* < 0.05) was used to evaluate the effect of TMS on local and global integration parameters.

#### Classification

For classification, we used the random forest implementation from the scikit-learn library (Pedregosa et al. 2011). As features, we included the nodal difference of *h* between the pre- and post-TMS data for all cortical nodes (N = 151; except class ‘undefined’) of the parcellation atlas by Power et al. (Power, Cohen et al. 2011). Critically, we used all cortical nodes of the atlas, not just nodes with significant changes in our prior analysis steps, thereby avoiding any bias in feature selection (Arbabshirani et al. 2017). The calculation of *h* was based on the individual co-classification matrices and the Power network assignments as module affiliation (Power, Cohen et al. 2011), thereby avoiding any leakage of information from the test to the training data. We evaluated the performance of the classifier using (i) a nested-cross validation (leaving out the two observations corresponding to one subject for testing) and (ii) an inner validation approach for the hyperparameter optimization of the random forest-classifier (using a sequential model-based optimization implemented by the Scikit-Optimize library; skopt, https://github.com/scikit-optimize/scikit-optimize), iteratively tuning the following parameters following the recommendations by Probst et al. (Probst, Wright and Boulesteix 2019): maximum depth of the tree, number of features, minimum number of samples and minimum number of samples required to be at a leaf node. We statistically validated the observed accuracy using permutation testing (*p* < 0.05, 5000 iterations) randomizing the class labels.

#### Parametric empirical Bayes (PEB) framework for DCM

The subject specific DCMs were taken to the second level where we used PEB routines for group level inference (Friston, Litvak et al. 2016); these routines assess how individual (within-subject) connections relate to group means, taking account of both the expected strength of each connection and the associated uncertainty. Specifically, we created three separate second-level PEB models to examine directional connectivity at baseline (preTMS) and changes after OCC-TMS and FRO-TMS within eight functional subnetworks. Next, we used Bayesian model reduction to test all nested models within each full PEB model (assuming that a different combination of connections could exist for each participant: Friston, Litvak et al. 2016) and to ‘prune’ connection parameters. The parameters of the best 256 pruned models were averaged and weighted by their evidence (Bayesian Model Averaging) to generate group estimates of connection parameters. Finally, we compared models using free energy and calculated the posterior probability for each model as a softmax function of the log Bayes factor. We report effects (connection strengths) as significant with a posterior probability > 0.95.

## Supplementary Figures

**Figure S1.**
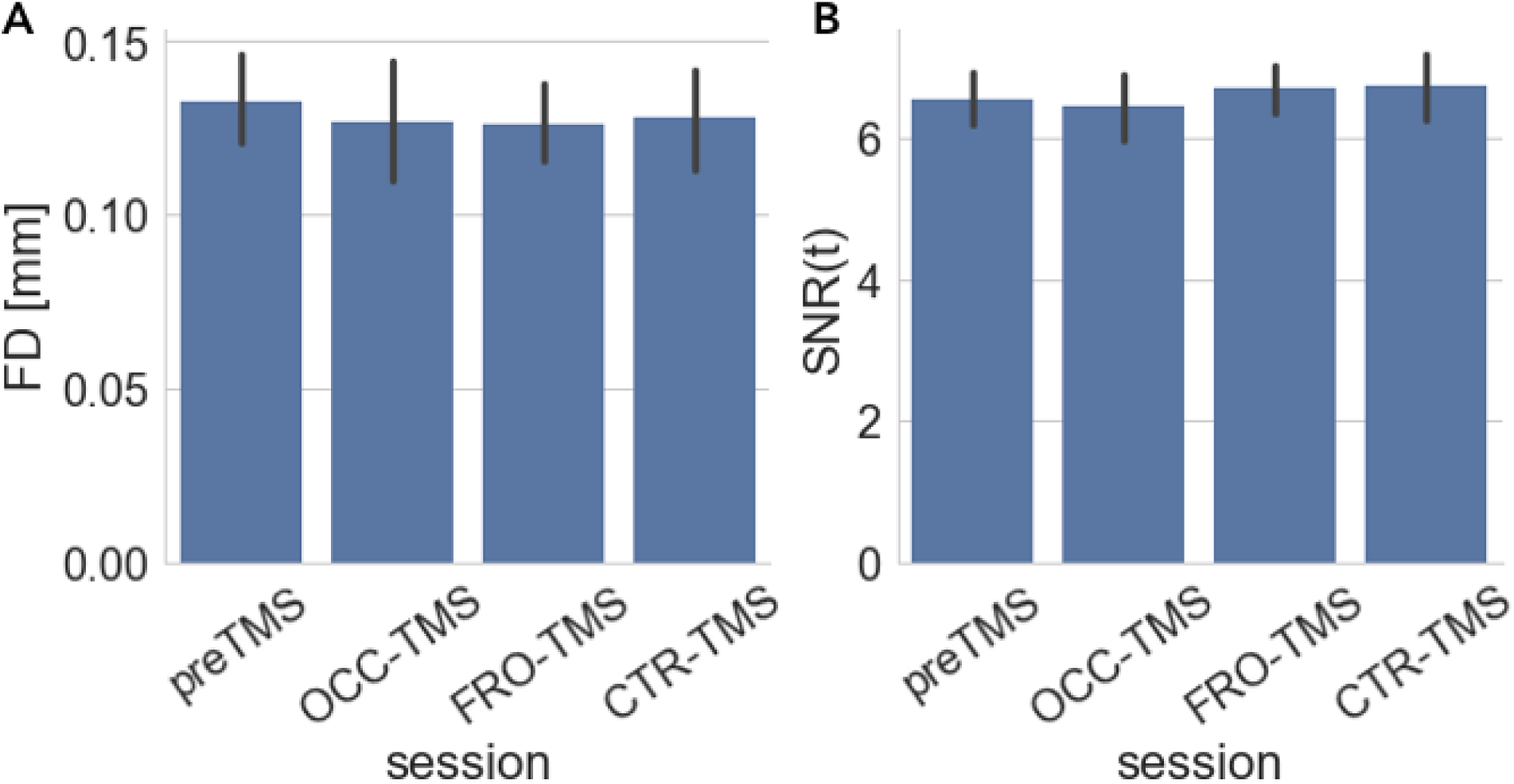
Image quality assessment. Bar plots represent (A) framewise displacement (FD) and (B) temporal SNR (SNR(*t*)) across TMS sessions. Error bars represent the 95% confidence interval of variance across subjects. Overall, there were not significant differences for any of the two parameters across TMS sessions (*p* > 0.05, repeated measures ANOVAs).

**Figure S2.**
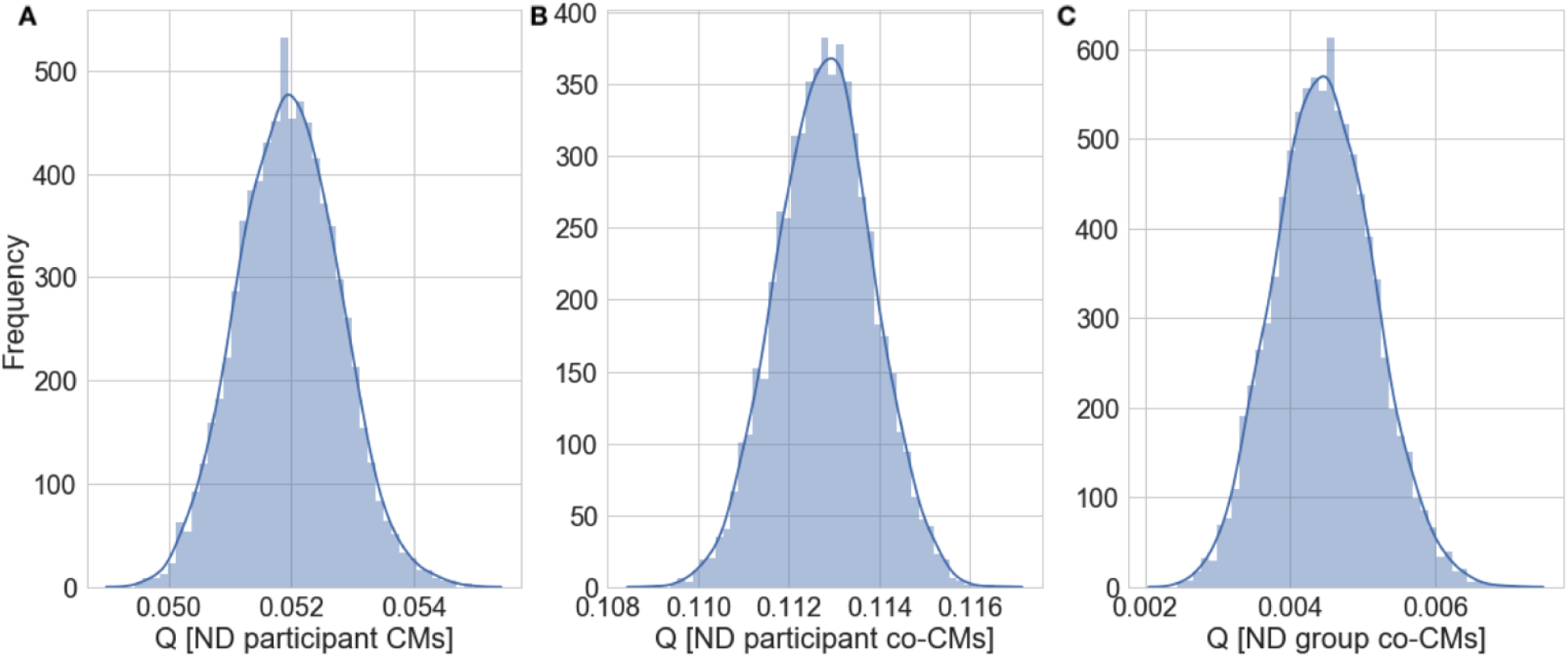
Statistical significance of the modularity results. Null model distributions (ND) of median Q values based on (A) the individual functional connectivity matrices, (B) the individual co-classification matrices, and (C) the group co-classification matrices. The modularity results were more modular than expected by chance when compared to the null model for the individual connectivity matrices (Q_pre_ = 0.181, Q_OCC_ = 0.143, Q_FRO_ = 0.165; p < 0.001, permutation testing), individual co-classification matrices (Q_pre_ = 0.585, Q_OCC_ = 0.544, Q_FRO_ = 0.588; *p* < 0.001, permutation testing) and group co-classification matrices (Q_pre_ = 0.266, Q_OCC_= 0.244, Q_FRO_ = 0.274; *p* < 0.001, permutation testing).

**Figure S3.**
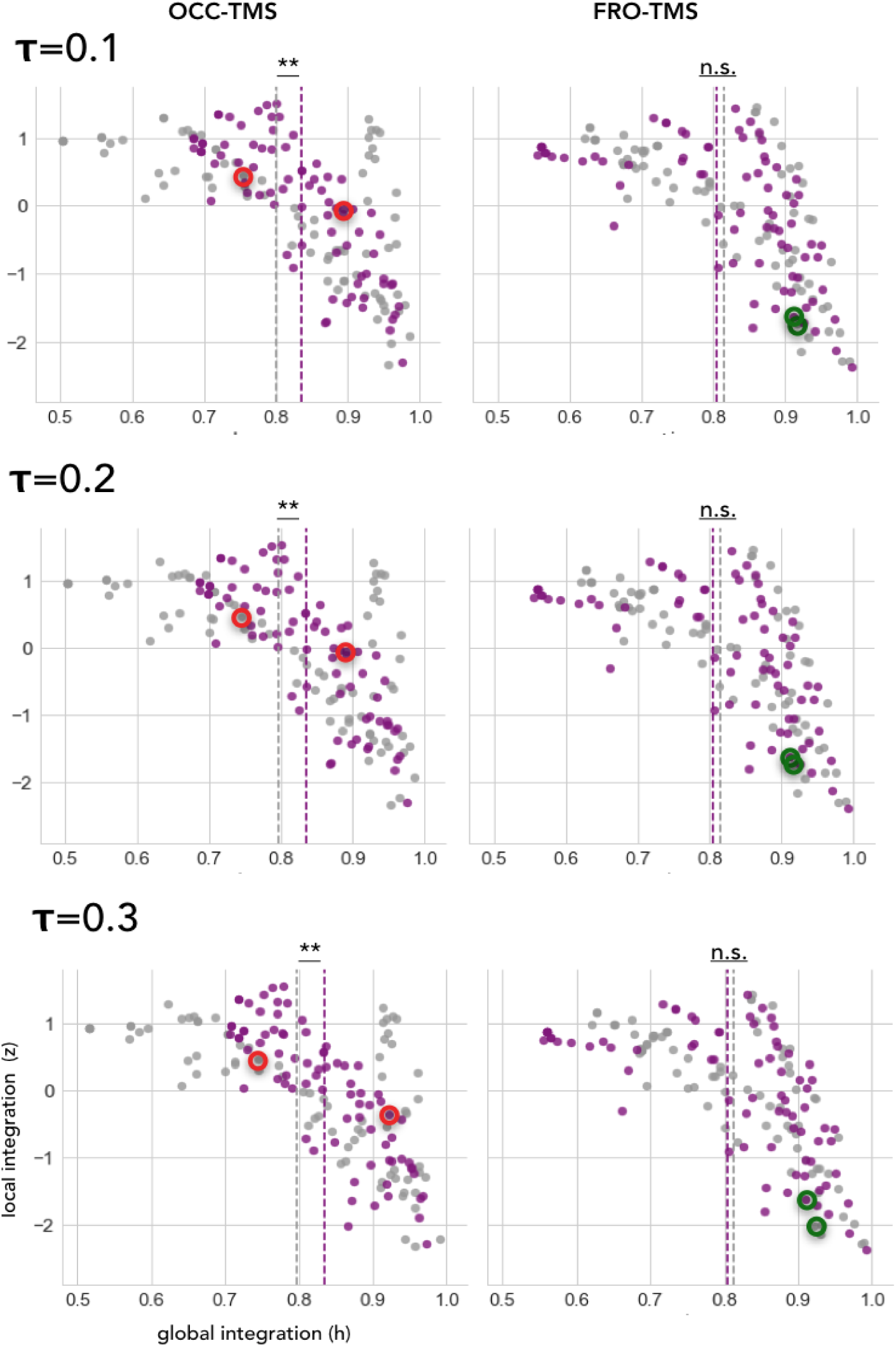
Effect of the parameter τ on the global functional integration results. Scatterplots of *z* vs. *h* before (grey) and after (violet) stimulation across the range of optimal values for the consensus modularity parameter *τ* (*τ* ≤ 0.4), which reproduces the global functional integration results obtained using the recommended value of *τ* = 0.4 (Lancichinetti and Fortunato 2012) in (Fig. 3C-D). ** *p* < 0.01, Wilcoxcon signed-rank test.

**Figure S4.**
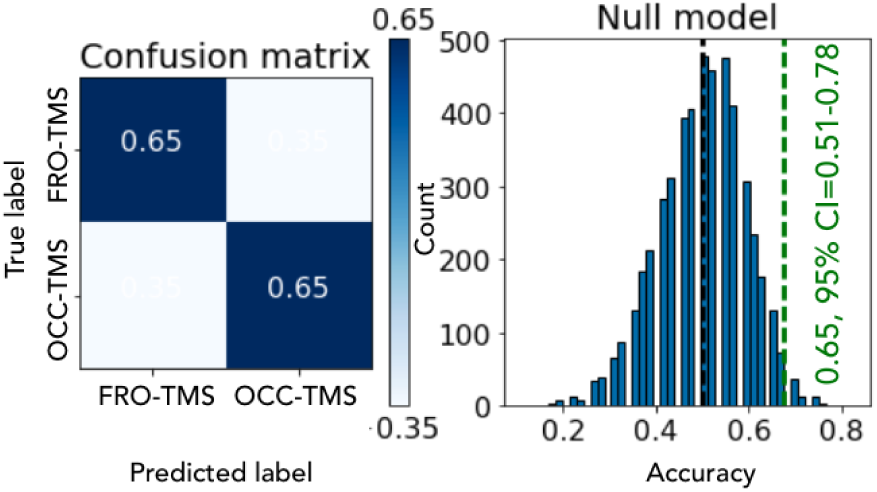
Classification results using linear SVM. Results of classification between VIS- and SAL-TMS data lead to a 65% prediction accuracy. (Left) Confusion matrix with the prediction accuracies for every class and (Right) null distribution of chance after 5000 permutations which shows that our results are significantly higher than chance (*p* = 0.029, permutation testing).

